# Universal features of Alternative splicing in response to diverse environmental stimuli in rice and the vital roles of TFs in AS regulation

**DOI:** 10.1101/2024.08.23.609440

**Authors:** Benze Xiao, Shuai Yang, Chengqi Wang, Fangyu Zhang, Yi Liu, Zhuowei Xiao, Guosheng Xie, Zhengfeng Zhang

## Abstract

**Background:** Pre-mRNA alternative splicing (AS) plays essential roles in response to environmental stimuli in plants. However, the universal and specific features of splicing in response to diverse environmental conditions remain not fully understood. Recent studies have shown the co-transcriptional characteristics of splicing, which lead to the reasonable speculation that the elements or factors regulating transcription can also affect splicing. Among of which, the effects of transcription factors on alternative splicing in plants under environmental stimuli are still confusing. A large amount 0f public available RNA sequencing data are valuable resources to be re-analyzed for answering questions beyond the aims of their original studies.

**Results:** We explored the universal features of AS using a standard RNA-seq dataset TENOR, which stems from rice samples under controlled diverse conditions to provide comprehensive and comparable AS analysis under various conditions. We found that AS widely occurs in rice under stimuli, with significant tissue specificity, temporal dynamics, commonality among different stresses or treatments as well as significant difference between differential alternative splicing and expressed genes (DASGs and DEGs) in rice under environmental stimuli. The majority of DASGs under various stresses are splicing factors and transcription factors. The correlation analysis shows that the expression level of transcription factors is significantly correlated with the PSI of AS events. The predominant transcription factors correlating with alternative splicing events come from bHLH, bzip and hsfa families. We validated the effects of transcription factors on AS by analyzing RNA-seq data from transcription factor mutants and found substantial differential AS events between mutants and wild type. Furthermore, the significant correlation was discovered between the transcription levels of transcription factors and splicing factors.

**Conclusion:** We found universal features of AS and the predominant AS events of SFs and TFs in plants under diverse environments. We propose that TFs might regulated AS of download genes partly by changing the patterns of their own transcription and splicing to further regulate the transcription of SFs. This work illuminate the studies on the possible mechanisms by which TFs modulate AS in plant, especially under environmental stimuli.

## Introduction

Global change in climate and environment is posing an increasing threat to crop growth and yield by bringing adverse factors including extreme temperature, water scarcity, salinization, flooding, soil contamination and so on (Renard & Tilman 2019; Li *et al*. 2022; Chen *et al*. 2024). As sessile organisms, plants have evolved systematic and exquisite programs to adapt or resist diverse environmental stresses. As the world’s rice production and grain quality have drastically declined due to the challenges posed by global climate change and abiotic stresses, it is crucial to develop rice cultivars maintaining or enhancing yield of rice under multiple abiotic challenges (Saud et al. 2022; Eckardt et al. 2023). To this end, one of the key measures is to explore the common and specific mechanisms in response to various environmental stimuli and discover the pleiotropic regulation factors (Sarma 2023).

More and more efforts have been put into the understanding of mechanisms of plant adaptation to multiple environmental stimuli (Shahzad et al. 2021). For example, it was reported that plants exhibit a notable up-regulation of acclimatin-associated metabolites in response to UV-B radiation and drought combined stress (Kaur et al. 2024). Transcription factors (TFs) MhDREB2A/MhZAT10 play a role in drought and cold stress response crosstalk in apple (Li 2023). Similarly, AtMYB68 play a crucial role in plant under the combined stress of heat and drought (Mingde Deng 2023).

AS is the process that can generate multiple distinct transcript isoforms from a pre-mRNA molecule through selecting different splicing sites (Petrillo 2023). It is suggested that pre-mRNA splicing variation is an important regulator in plant adaptation to environmental stresses, such as heat, drought, cold, mineral deficiency and so on (James et al. 2012; Dong et al. 2018; Zhang & Xiao 2018; Ling et al. 2021; Ye et al. 2023). AS also plays vital regulation roles in hormone responses in plants. For instance, Abscisic acid (ABA) can effectively control plant growth or improve stress resistance not only by regulating TFs expression, also via mediating alternative splicing of TFs (X et al. 2022). Though AS was extensively studied in plants in response to environmental stimulus, the comparable studies on AS response to multi stimuli and the identification of pleiotropic factors regulating AS in plant adaptation to complex stress conditions are still lacking.

Splicing factors (SFs) are well known trans factors to regulate AS (Reddy et al.). For example, SCR106 or SmEb can modulate abiotic stress responses by maintaining the RNA splicing in rice (Yang *et al*. 2022; A *et al*. 2023). Additionally, recent studies discovered the actuality that splicing occurs co-trancriptionally, suggesting the relevance of RNApolII, transcription factors and chromatin state to alternative splicing. For example, a recent research in human reported that TFs are involved in the regulation of intron retention by comparing regions of open chromatin in retained and excised introns (Ullah *et al*. 2018; Jabre *et al*. 2019; Malla *et al*. 2022; Ullah *et al*. 2023). A zinc finger protein 207 (ZFP207) was demnostrated to play a pivotal role in mouse embryonic stem cells (ESC) maintenance not by transcriptionally regulating relevant genes but via controlling AS networks through acting as an RBP (Malla et al. 2022). Similar studies have been reported in plants (Jabre et al. 2019). An Arabidopsis SERRATE (SE) gene, encoding a Zinc-finger protein, played key function in plant response to salt stress by regulating pre-mRNA splicing of several well-characterized marker genes associated with salt stress tolerance (Yang et al. 2022). However, the regulation roles of transcription relevant factors on AS in plants is yet far from clarity. The function of trancription factors on AS in response to environmental stimuli is still unrevealed in plants. the comprehensive relation between transcription factors and AS remained rarely reported in rice under different environmental stimuli.

The increasing number of high confidant and throughput RNA sequencing data have provided unprecedented chance to explore AS events. In the present study, we employed the public available data to explore AS response to environment conditions and the regulation function of TFs on AS. A time-course RNA-seq dataset (TENOR) was provided on rice under 10 abiotic stress conditions and two plant hormone treatment conditions available on Sequence Read Archive (SRA) in NCBI with accession number DRP000997 (Kawahara et al. 2016). This dataset provided unique advantages such as all experiments performed at a single laboratory under standardized conditions to better under the actual similarities and differences among various growth conditions. However, these valuable data was only analyzed at gene expression levels, leaving AS response to various environments unexplored. In this study, we will use data from TENOR to comprehensively analyze the universal AS features in rice under multiple growth conditions. We found the wide involvement of TF in AS changing after abiotic stresses and further validate the effects of TFs on AS using the RNA-seq data from a few TF mutants under stress and explore the regulation roles of TF in AS by the transcription relationship of TFs with SFs.

## Results

### AS events are frequent under various environmental stimuli

To comprehensively compare the AS landscape in rice under different environmental conditions, we analyze publicly available RNA-seq data, TENOR from a previous time-course study in rice under 10 conditions, including cadmimum, cold, flood, drought and osmotic stresses and JA and ABA hormone treatments (Table S1), following the framework mentioned in Figure 1 (Kawahara *et al*. 2016).

**Figure 1.**
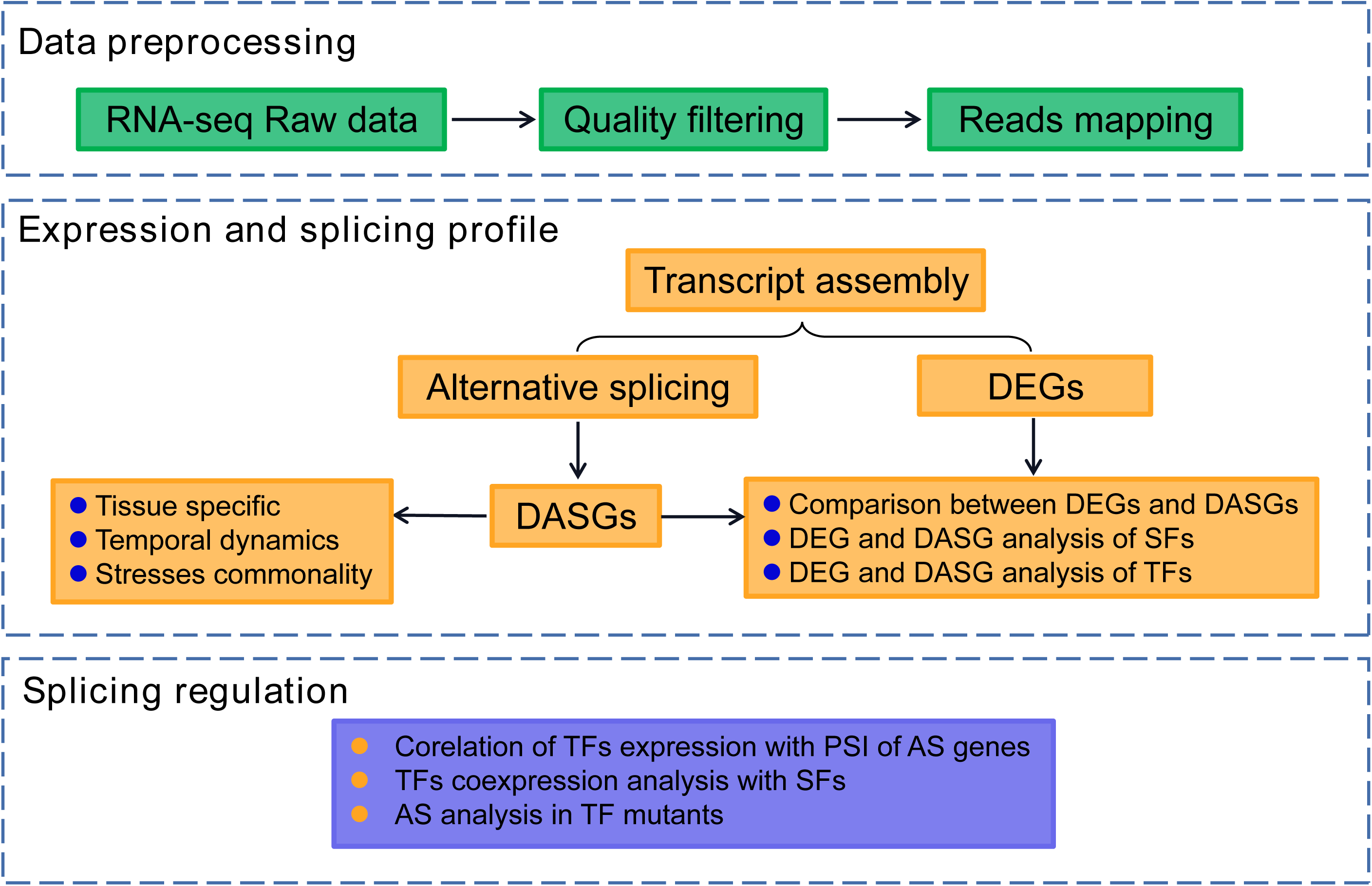
Overview of workflow used in this study. The workflow includes data preprocessing, differential gene expression and alternative splicing analysis and the regulation of alternative splicing.

A total of 238 RNA-seq samples were collected from TENOR. After filtering out low quality reads, 2234 million reads were obtained and 88.11% of them were mapped on the reference genome. Through transcripts assembly for individual samples and integration of transcripts in all samples, we finally obtained 35,952 genes and 64,681 transcripts, 24,659 multi-exon genes and 48,592 multi-exon transcripts. Among them, the number of transcripts with less than 10 introns accounted for approximately 84.2% (Figure S1), which is close to the 87.9% reported by Dong et al. (2018), indicating that the high accuracy of the transcript annotation. The median and mean intron sizes were 161 and 465 bp, respectively.

Extensive AS events were detected in rice root and shoot under various environmental conditions. The ratio of genes exhibiting AS in the expressed genes was ranging from 14.36% to 24.36%, referring to 2,902 AS genes in 20,213 expressed genes in root under JA treatment at 12h and 5,342 AS genes in 21,932 expressed genes in root under cold stress at 12h. The total number of genes undergoing AS events was 13,025, accounted for ∼52.8% of the total 24,659 expressed intron containing genes. This results indicated the extensive AS events in rice under environmental stimuli (Figure S2). Five main types of AS evens, including intron retention (IR), alternative 3’ splice sites (A3SS), alternative 5’ splice sites (A5SS), exon skipping (ES), and mutually exclusive exons (MXE) were detected in rice. Among of which, IR is the predominant pattern and MXE is the least AS type in all the environmental conditions (Figure 2).

**Figure 2.**
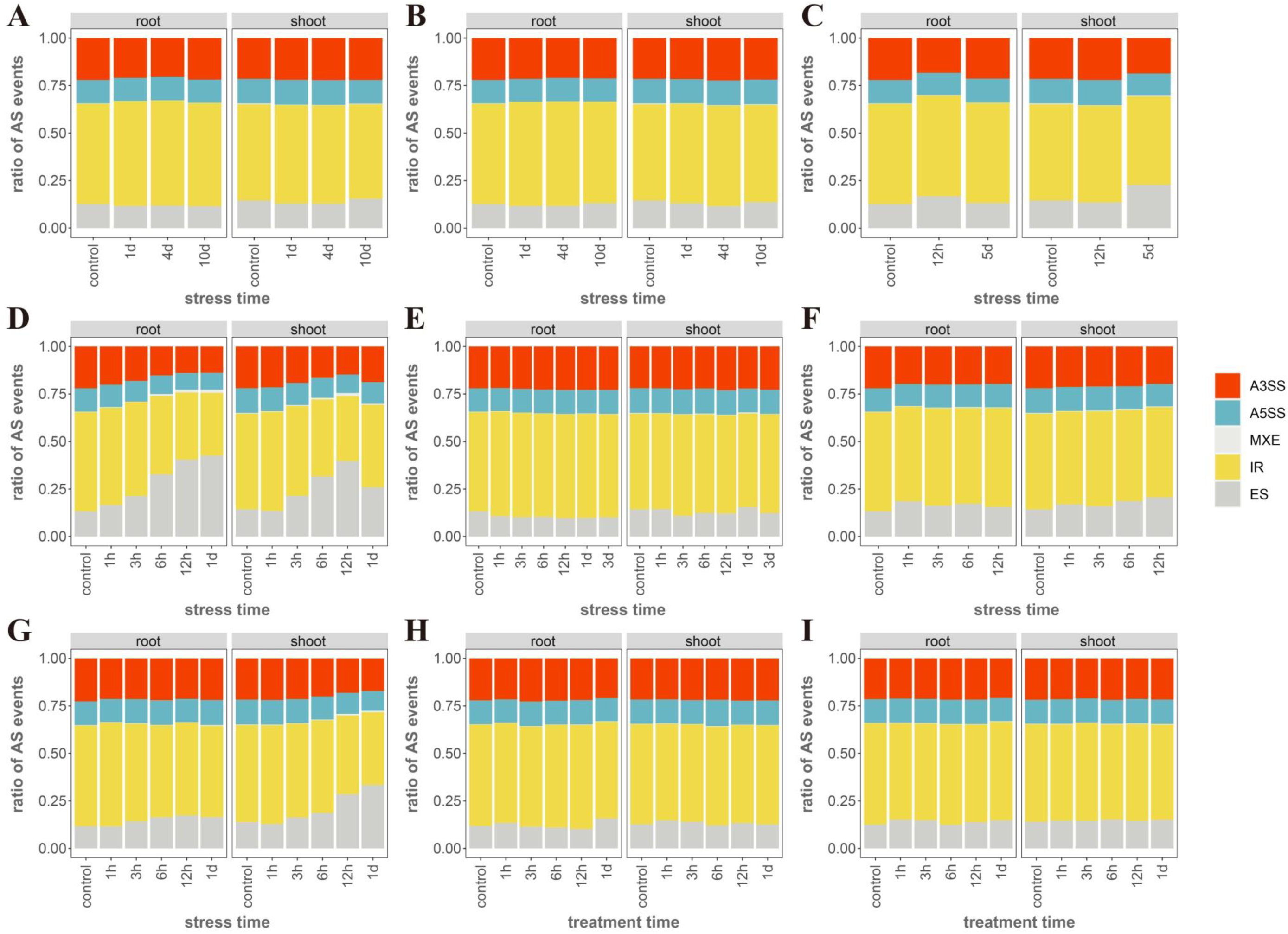
The ratio of five main AS types in response to various environmental stimuli. We computed the ratio of different AS types in shoots and roots, respectively. A. Very low Cd stress. B. Low Cd stress. C. High Cd stress. D. Cold stress. E. Flood stress. F. Osmotic stress. G. Drought stress. H. JA treatment. I. ABA treatment.

### Environmental stimuli induced differential AS genes are tissue specific

To unveil if environment induced different AS events are tissue specific, we compared the differential AS genes (DASGs) induced by various stress or treatments. For a specific stress, we first selected genes with ΔPSI≥10% in any AS event between any treat time point and control condition as DASGs, respectively in shoot and root (Table S2-S15). The DASGs subsequently were compared between root and shoot under each condition. In general, we found a small faction of DASGs were shared between root and shoot. The lowest and the highest number of shared DASGs between root and shoot were 84 and 620, which were under very low concentration of cadmium and cold treatments and occupied 11.5% and 28.8%, respectively (Figure 3). To further explore the AS responsive in different tissues under environmental stimuli, GO annotation was performed for DASGs in shoot and root, respectively. The results show GO terms of DASGs under diverse conditions were overall not consistent between these two tissues (Figure 4, S3). Some shared GO terms are unexceptionally enriched in RNA binding, mRNA splicing, nucleic acid binding, splicesome, etc. For tissue specific DASGs, the significant GO terms include signal transduction, oxidase activity, dephosphorylation, ion transport and so on (Figure 4, S3). These results indicate the stresses or hormones induced DASGs show distinct tissue specific manner. Meanwhile, RNA binding and splicing related functions and mechanisms were similar among tissues under external stimuli.

**Figure 3.**
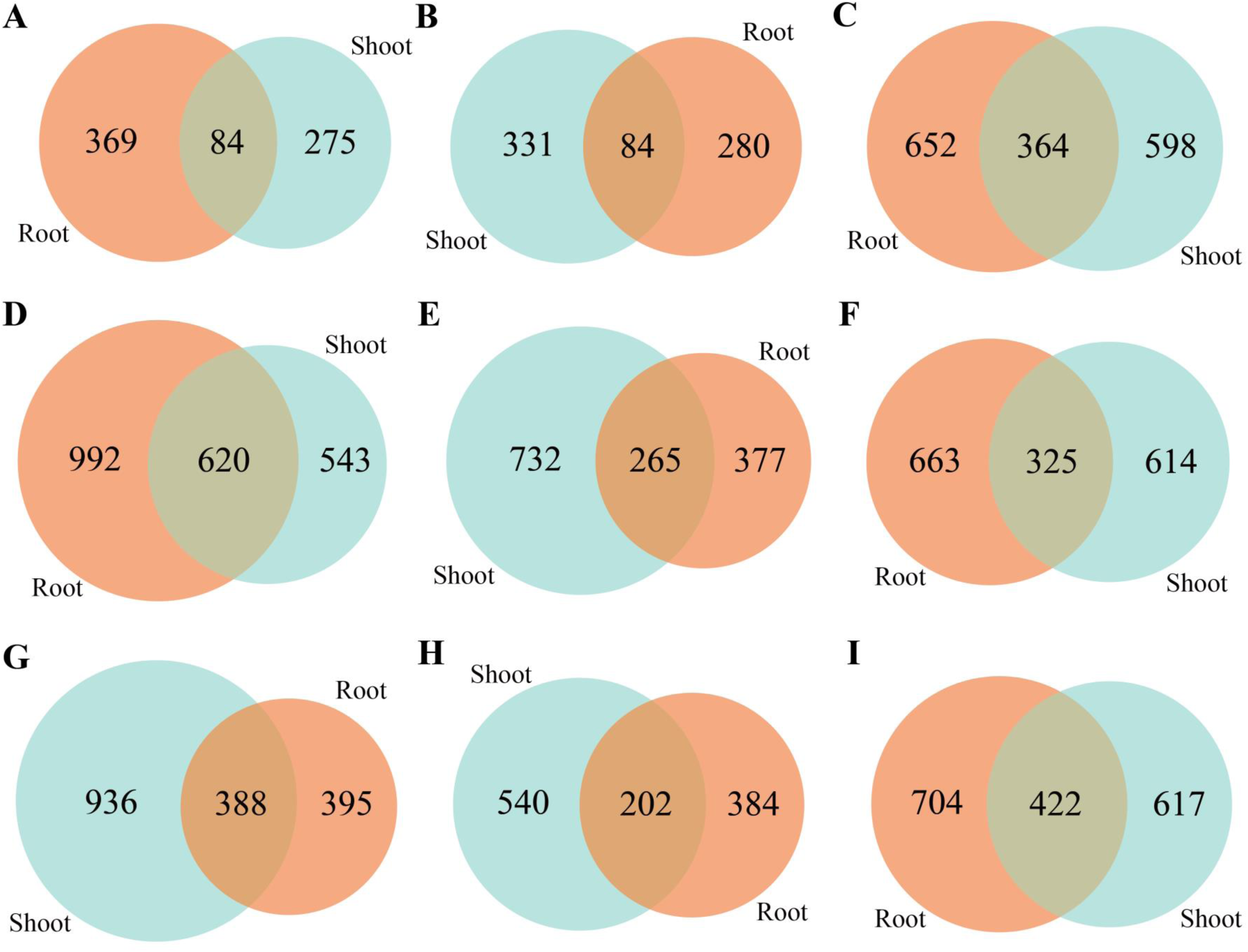
Tissue specific of DASG. DASGs were screened based on ΔPSI between each treatment time and corresponding control. DASGs at all timepoints were combined for each treatment in shoots and roots, respectively. The venn diagrams indicate overlaps of DASGs in shoots and roots under each treatment. A. Very low Cd stress. B. Low Cd stress. C. High Cd stress. D. Cold stress. E. Flood stress. F. Osmotic stress. G. Drought stress. H. JA treatment. I. ABA treatment.

**Figure 4.**
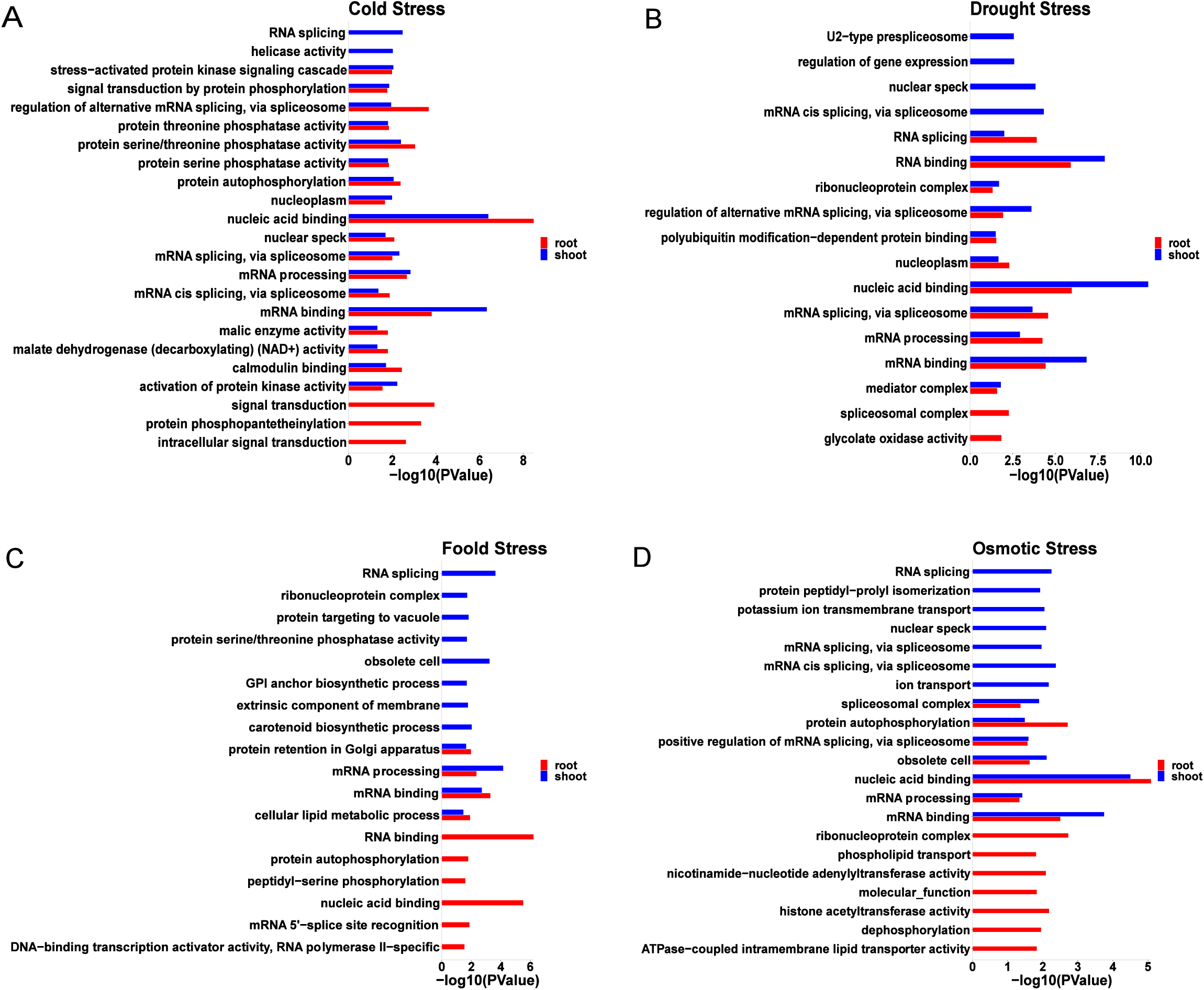
Functional GO analysis of DASGs in rice shoots and roots. Graphs show functional categorization (GO) of DASGs in shoots (blue) and roots (red) under cold (A), drought (B), flood (C) or osmotic (D) stress. The log10(1/P) values represents the enrichment ratio of each GO terms.

### Environmental stimuli induced DASGs are fluctuated with the processing duration

To test whether the AS pattern changes over time in response to external stimuli, we examined DASGs at different time points under each condition. It was found that only a small proportion of genes exhibiting DAS throughout the whole treatment process (Fig. 5). For example, a total of 1,161 genes were identified showing differential AS in 5 time periods in shoots after cold stress. Among of them, AS changing are induced in 569 genes (49%) at only one time point after low temperature stress, whereas only 20 DASGs (1.72%) maintained along with the entire cold processing period. The number of DASGs increased first and then decreased, and reached the peak at 12 h after cold stress in rice shoots (Fig. 5A). The similar situation occurs under cold stress in rice roots, where a total of 1,606 DASGs were detected upon cold stress, including 737 (45.9%) DASGs at one time point after stress and only 123 (7.66%) DASGs in response to cold stress along the entire stress process. The number of DASGs increased with the stress continuously, and reached the peak at 24h after cold stress in rice roots (Fig. 5B). The feature of mRNA splicing fluctuation with stress processing occures similarly in other conditions, including drought (Fig. 5C for shoot and 5D for root), flooding (Fig. S4A for shoot and S4B for root), osmotic (Fig. S4C for shoot and S4D for root), Cd stress (Fig. S5A-C for shoot and S5D-F for root), JA (Fig. S6A for shoot and S6B for root) and ABA treatments(Fig. S6C for shoot and S6D for root). These results indicated that AS responds to environmental stimuli in plants in a temporal dynamic manner.

**Figure 5.**
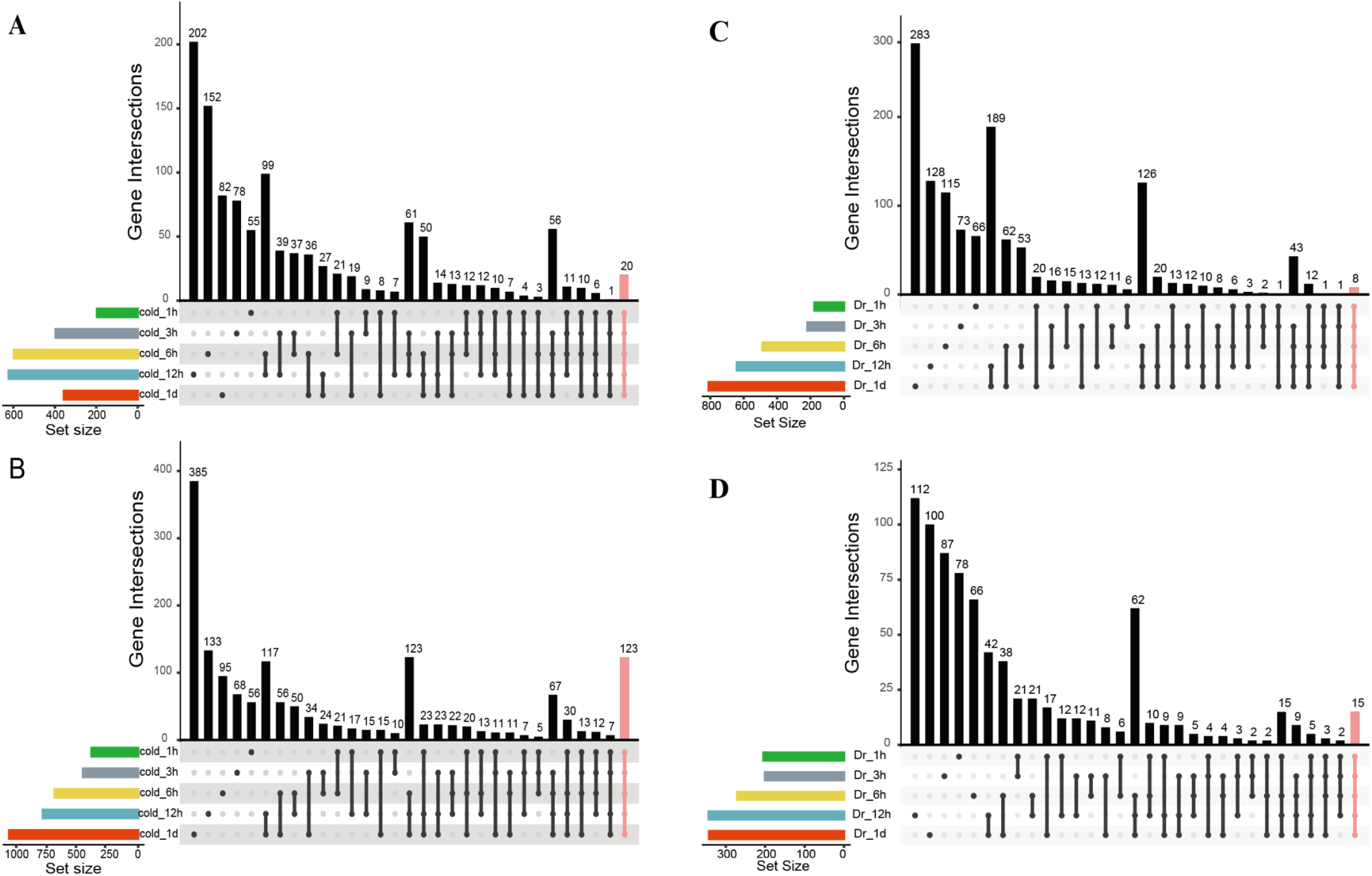
The temporal fluctuation of stress-induced DASGs. The DASGs were dynamically changing with the treatment duration under environmental stimuli. The time specific and shared DASGs are shown for shoots (A) and roots (B) under cold stress or for shoots (C) and roots (D) under drought stress.

### Some DASGs are shared in responsive to different environmental conditions

To detect if there are overlapped DASGs in responding to different stresses, we compared the DASGs among different conditions for shoots and roots, respectively. The results showed that there were indeed overlapped DASGs among various environmental stimuli. The most noteworthy is that out of 2338 DASGs in shoots (Fig. 6C) and 2282 in roots (Fig. 6D) under cold, drought, flood and osmotic stress, about 53% and 46% were concurrently responding to 2 to 4 different abiotic stresses, respectively. About 9.67% DASGs were regulated simultaneously by all 4 stresses in shoots. 16.89% and 26.39% DASGs respond to 3 and 2 abiotic stresses in shoots, respectively (Figure 6C). Similarly, 7.49%, 15.16% and 23.58% DASGs in roots simultaneously respond to 4, 3 and 2 abiotic stresses, respectively (Figure 6D). Similarly, there were 115, 145, and 648 DASGs in shoots and 120, 91, and 634 DASGs in roots under extremely low, low, and high concentrations of cadmium stress, respectively. Among of them, 88 and 124 genes underwent differential AS simultaneously under 2-3 cadmium concentrations of stress (Figure 6A, B). Besides these stresses, a set of DASGs were simultaneously regulated by different hormone treatments. About 24.11% and 20.14% DASGs were overlapped between ABA and JA treatments in shoots (Figure 6E) and roots (Figure 6F), respectively. All these results show that some genes might play pleiotropic roles in plants responding to various stresses or hormone treatments. These genes are worth to undergo in-depth research to promote the cultivation of varieties with multi-tolerances to diverse adverse conditions.

**Figure 6.**
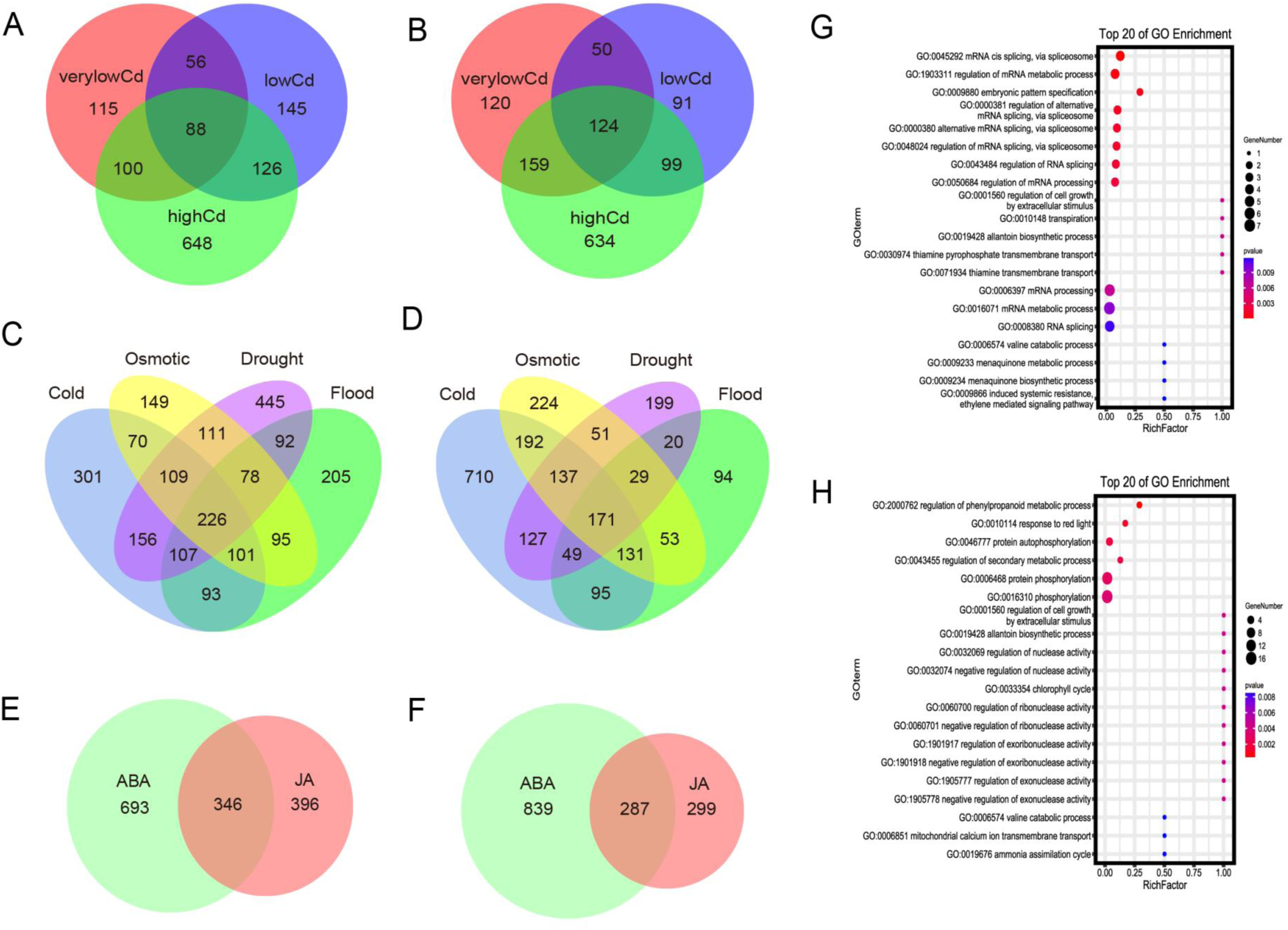
Shared DASGs among various environmental stimuli. The venn diagrams show the overlap of DASGs in different environmental conditions. The DASGs intersection among different concentration of Cd stress were shown for shoots (A) and roots (B). The DASGs intersection among cold, drought, flood and osmotic stresses were shown for shoots (C) and roots (D). The DASGs intersection among ABA and JA treatments were shown for shoots (E) and roots (F). The enrichment GO terms for shared DASGs under cold, drought, flood and osmotic stresses are shown for shoots (G) or roots (H).

We further conducted GO annotation for overlapped DASGs in responding to different combinations of stresses. The top enriched functions for DASGs overlapped in responding to cold, drought, flood and osmotic stresses are relevant to mRNA splicing or processing GO terms in either shoot (Fig. 6G) or root (Fig. 6H). The consistent results were detected for GO annotation of DASGs overlapped in other conditions (Fig. S7). These results indicated the significant roles of mRNA splicing regulating genes in plants coping with multi environmental stimuli.

### Stress-induced DEGs and DASGs influence different biological processes

Since gene transcriptional level changing (differential expressed genes, DEGs) is the main focus for most studies on plant stress response, we are curious to inspect the intersection between DEGs and DASGs. We generally found only a small faction of DASGs were overlapped with DEGs in responding to various conditions. Particularly under cold stress, the overlapped gene number between DASG and DEG was 4, 90, 163, 292, and 41, only occupying 2.04-46.95% of total DASGs in shoots under 1 hour, 3 hours, 6 hours, 12 hours, and 1 day duration of stress (Figure 7A). In roots under cold stress, 82, 87, 186, 234, and 340 overlapped genes were identified between DEGs and DASGs at 5 time points after stress. The ratio of DASGs overlapped with DEGs to the total DASGs was 21.41-32.14% (Figure 7B). The similar pattern was shown under drought (Figure 7C and D), flooding (Figure S8A), osmotic (Figure S8B) and Cd stresses (Figure S9) and JA (Fig. S10A and B) and ABA treatments (Fig. S10C and D). We further check the extent of the function overlaps of DEGs and DASGs. The results in cold stress show except one GO term, metal ion transport was enriched in both DASGs and DEGs, no other enriched annotation were overlapped. DASGs specific are mainly enriched in mRNA binding and splicing GO terms. DEGs specific show an extensive enriched GO terms, including ion transport, response to stress, regulation of defense response, mitochondrial respiratory chain complex, signal transduction and so on (Figure 7E, F). No significant overlapped GO terms was enriched for both DEGs and DASGs under drought stress (Figure 7G, H). Similar results were found under other environmental conditions in the present study (Figure S11-12) and indicated that stress-induced DEGs and DASGs influence different biological processes. Therefore, it will lose important genes and clues if we only pay attention to transcription level regulation without post-transcription processing, such as AS, regarding to plants responding to environment stimuli.

**Figure 7.**
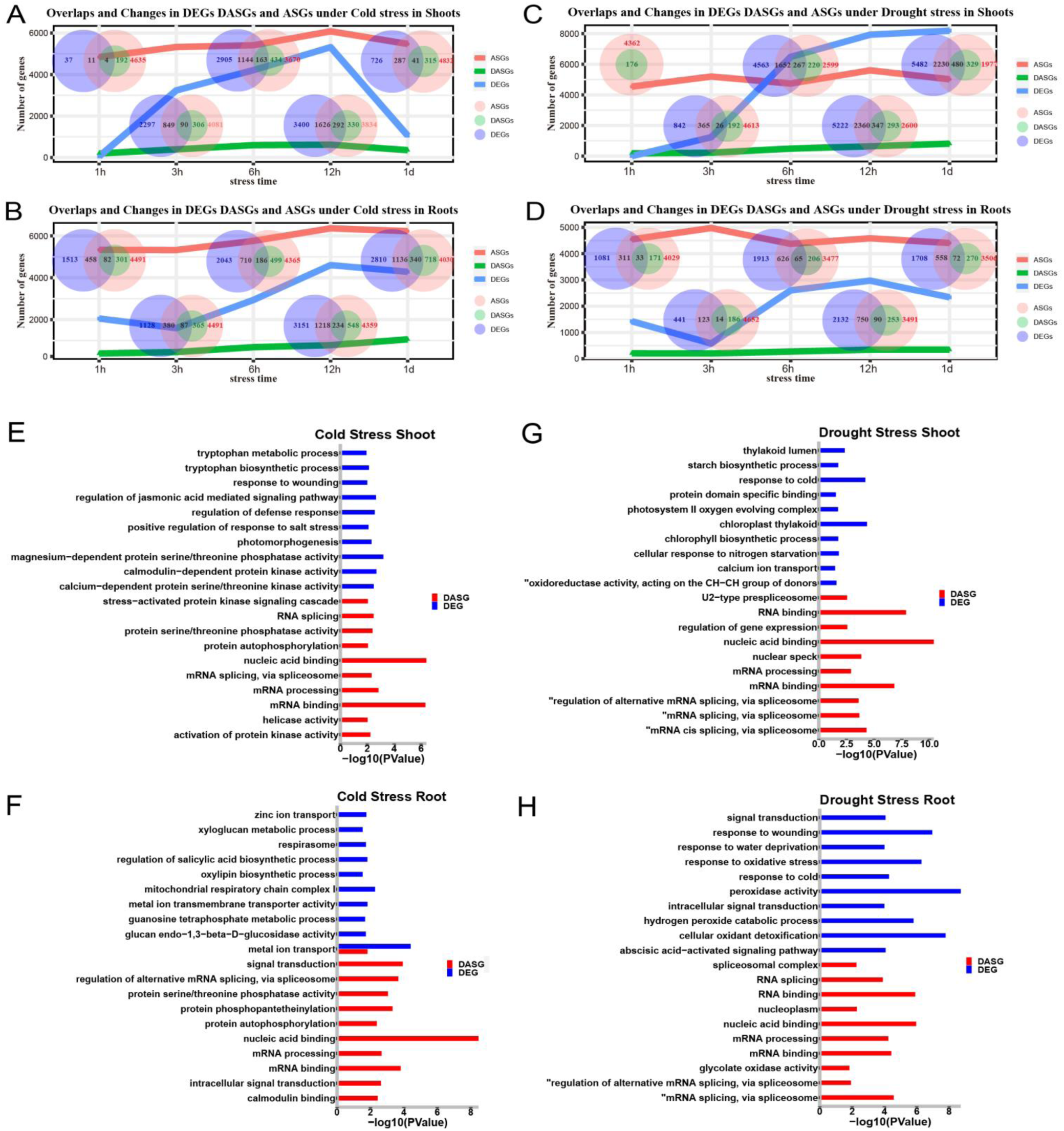
Comparison and GO annotation of DEGs and DASGs under cold or drought stress. (A-D) The line charts and venn diagrams represent the number and overlap of DEGs, ASGs and DASGs in shoots and roots under different cold or drought stress durations. (E-H) The enrichment GO terms for DASGs (red) and DEGs (blue) in shoots and roots under cold or drought stress.

### Splicing factors undergoing frequently alternative splicing in response to abiotic stresses

The above results show that a large amount of DASGs responding to various environmental stimuli were enriched in GO terms such as mRNA binding, splicing factors, and splicing regulation, no matter which stress treatment or tissue. It was speculated that splicing machine related genes plays a crucial role in plants responsive to stress. They probable regulate splicing patterns of other downstream stress responsive genes through firstly modulating the transcription and splicing of themselves. Serine/arginine-rich splicing factors (SFs) family play important roles in the assembly of spliceosome and the regulation of pre-mRNA splicing. Barta et al. (2010) identified a total of 22 SFs in 6 subfamilies including SR, SCL, SC, RSZ, RS2Z, and RS in rice. In order to systematically explore the involvement of splicing machine themselves in stress response, we conducted a comprehensive analysis of SF genes in environmental stimuli from both transcription and alternative splicing levels.

Indeed, most SF coding genes show relative high level of transcription under various abiotic stresses and plant hormone treatments, except SCL28 (Os03g0363800), SC25 (Os03g0388000) and RS2Z39 (Os05g0162600) expressed at are relatively lower level unpon all conditions (Figure 8A). Among of them, RSZ23 (Os02g0610600) exhibit persistent high expression level under all conditions in both tissues. SR33 (Os07g0673500) and SR32(Os03g0344100) show high level of transcription under all conditions in shoots. Additionally, the expression levels of SR33 and SR32 in shoots decreased with the prolongation of stress time in multiple treatments such as cadmium, low temperature, drought, JA, and ABA. The results indicated the importance of SFs in plants responding to environmental treatments (Figure 8A). Furthermore, SFs function in a tissue and treatment time specific manner. Intriguingly, although several SFs within each of the six subfamilies share similar protein domains, there is no clear pattern or similarity in their expression levels, indicating that SFs from the same subfamily do not have functional redundancy and may perform unique functions in pre-mRNA splicing regulatory network.

**Figure 8.**
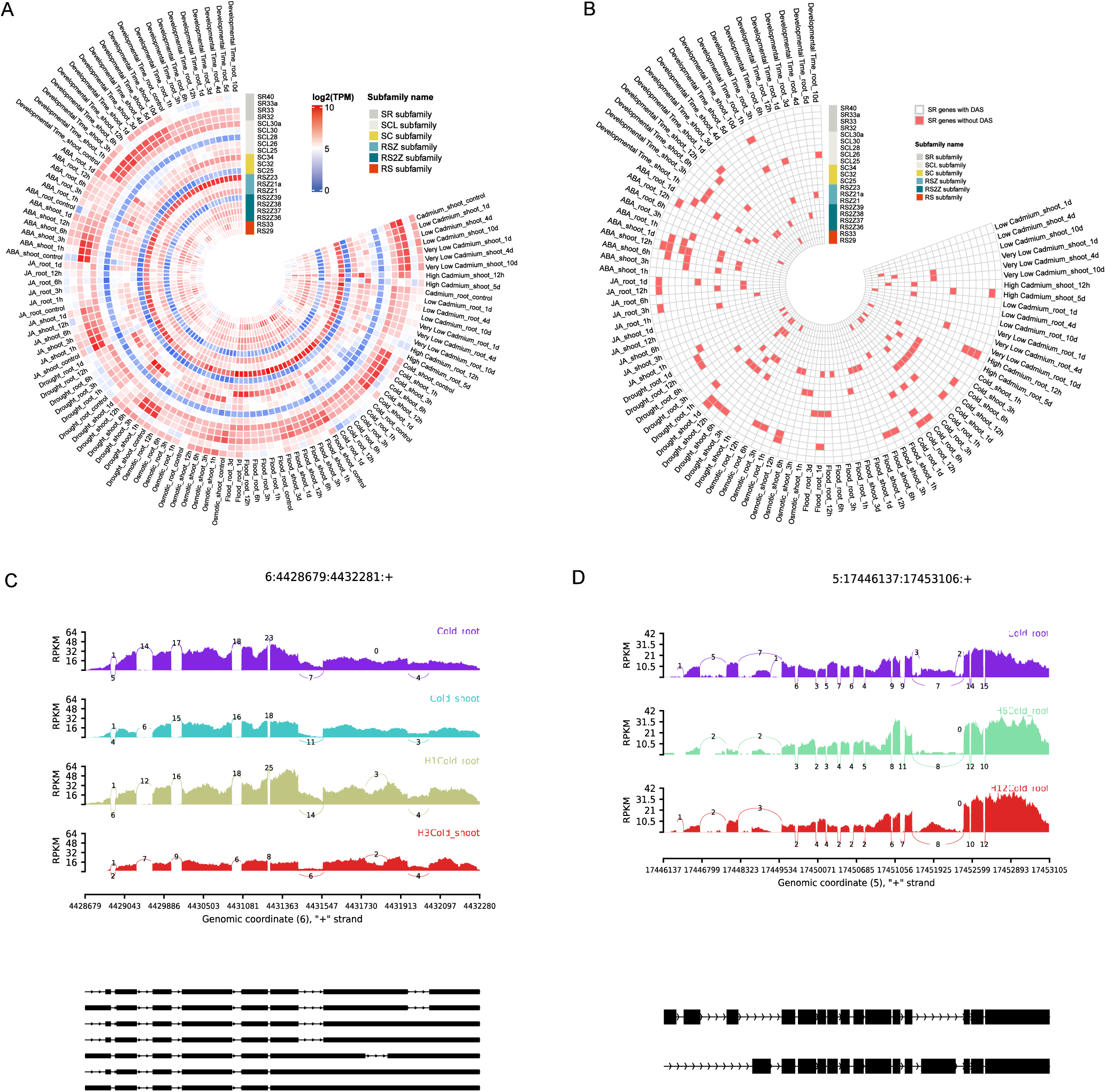
Expression and AS profiles of splicing factors under environmental stimuli. Circle diagrams show the transcriptional levels (A) and DASGs (B) of splicing factors coding genes in shoots and roots under various conditions. The representative DAS events of SF coding genes are shown for an intron retention event of gene RSZ21a (C) and an exon skipping even of gene SR33a (D) under cold stress.

It is notable that except modulating transcription levels, SFs form different splicing isoforms when responding to environmental stimuli. Most SFs undergo differential alternative splicing at multiple time points under multiple stimuli (Figure 8B). Among a total of 15 SFs that underwent differential AS events, 12 of them showed differential splicing pattern under two or more stress conditions. For example, *SC33a* undergoes differential AS under ABA treatment and cold, drough, flood and Cd stresses. *RSZ21a* (*Os06g0187900*) undergoes differential alternative splicing under both Cd, ABA and cold, drough, flood and osmotic treatments. The representative AS events were shown for intron retention in gene RSZ21a (Figure 8C) and exon skipping in gene SR33a under cold stress (Figure 8D). The results indicated the extensively roles of SR factors in plants responding to environmental treatments at the splicing level with a condition specific manner.

### The potential roles of TFs in AS regulation in responsive to abiotic stresses

To decipher if transcription factors has an impact on the AS regulation in rice under environmental stimuli, we firstly examined the response to environmental stimuli of TFs at both transcriptional and AS level. The results show that some TFs respond to environments specifically at transcription level, whereas other TFs respond at AS level or at both transcription and splicing levels (Figure 9), indicating the importance of AS of TFs in environmental responses in plants. We totally identified 158 TFs undergoing differentially expressed and differentially AS upon environmental stimuli, including multiple families such as ARF, bHLH, bZIP, C2H2, GATA, HSF, MYB, ERF, NAC, WRKY and so on (Figure 9, Table S16). Some of these TFs have been well demonstrated to function in plants adaptation to abiotic stresses. The emerging evidences demonstrated that pre-mRNA splicing is a co-transcriptional process and to some extent relevant to TFs. These clues indicate the crucial roles of TFs at AS level in response to environments. To further explore the possible relevant of TFs with AS regulation under environmental stimuli, we subsequently evaluated the relationship of transcription levels of TF with PSI of splicing events in other genes in rice under cold, drought and flood stresses at time series using Pearson correlation test (r>0.5 and p<0.05). The correlation network shows apparent and extensive correlation between TFs transcripition level and PSI of other gene splicing events (Figure 10). Take the drought stress as example, 41 members in bHLH TF family are significantly associated with alternative splicing events (Figure 10D). For instance, *Os08g0432800* and *Os03g0741100* is significantly correlated with 77 and 43 AS events, respectively. 40 members in bZIP family and 14 members in Hsfa family are significantly correlated with AS events, respectively (Figure 10E, F). For example, *Os07g0644100* and *Os06g0622700* in bZIP family are closed related to 67 and 46 AS events, respectively. *Os03g0795900* and *Os03g0161900* in Hsfa family closed related to 52 and 43 AS events, respectively. The profound correlation of TF transcription and AS events was widespread in other environmental conditions. The relationship of bHLH, bZIP and Hsfa with AS events were displayed under cold (Figure 10 A-C), drought (Figure 10 D-F) and flood stresses (Figure 10 G-I), respectively. These results demonstrated the potential regulatory function of TF in AS, beyond the regulation of TFs on gene transcription level.

**Figure 9.**
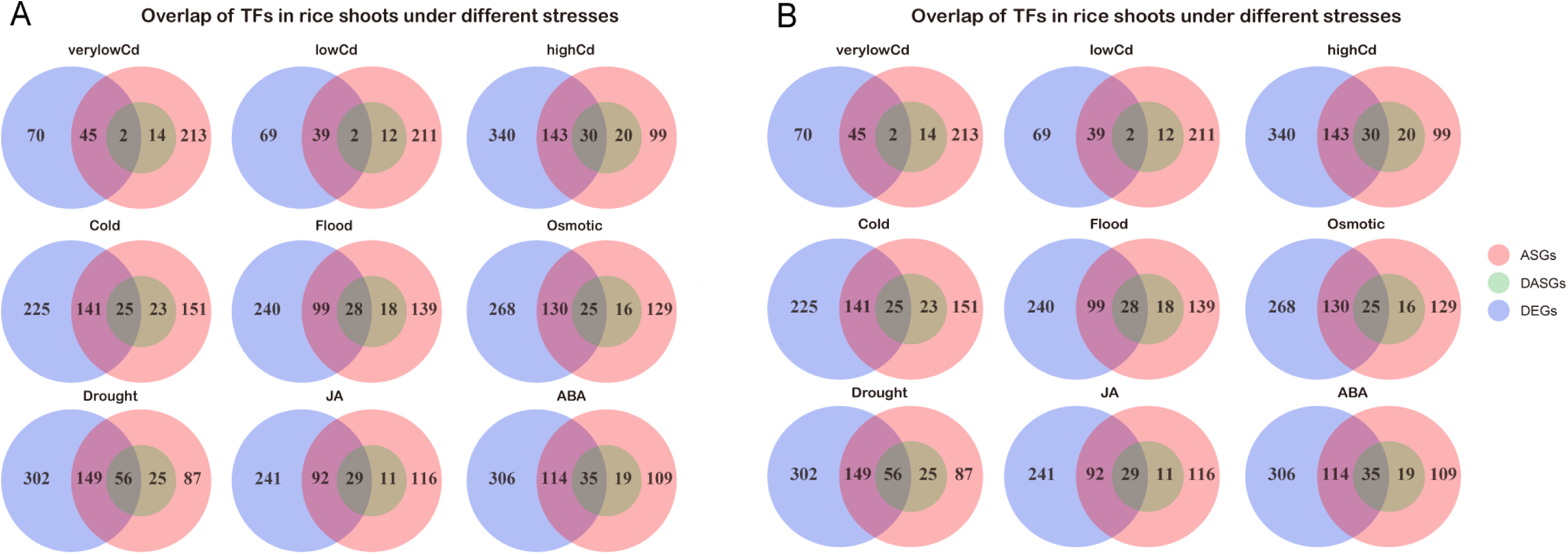
Comparison of differentially expressed and alternative splicing TFs. The venn diagrams show the comparison of TFs ASGs, DASGs and DEGs in shoots (A) and roots (B) under different conditions.

**Figure 10.**
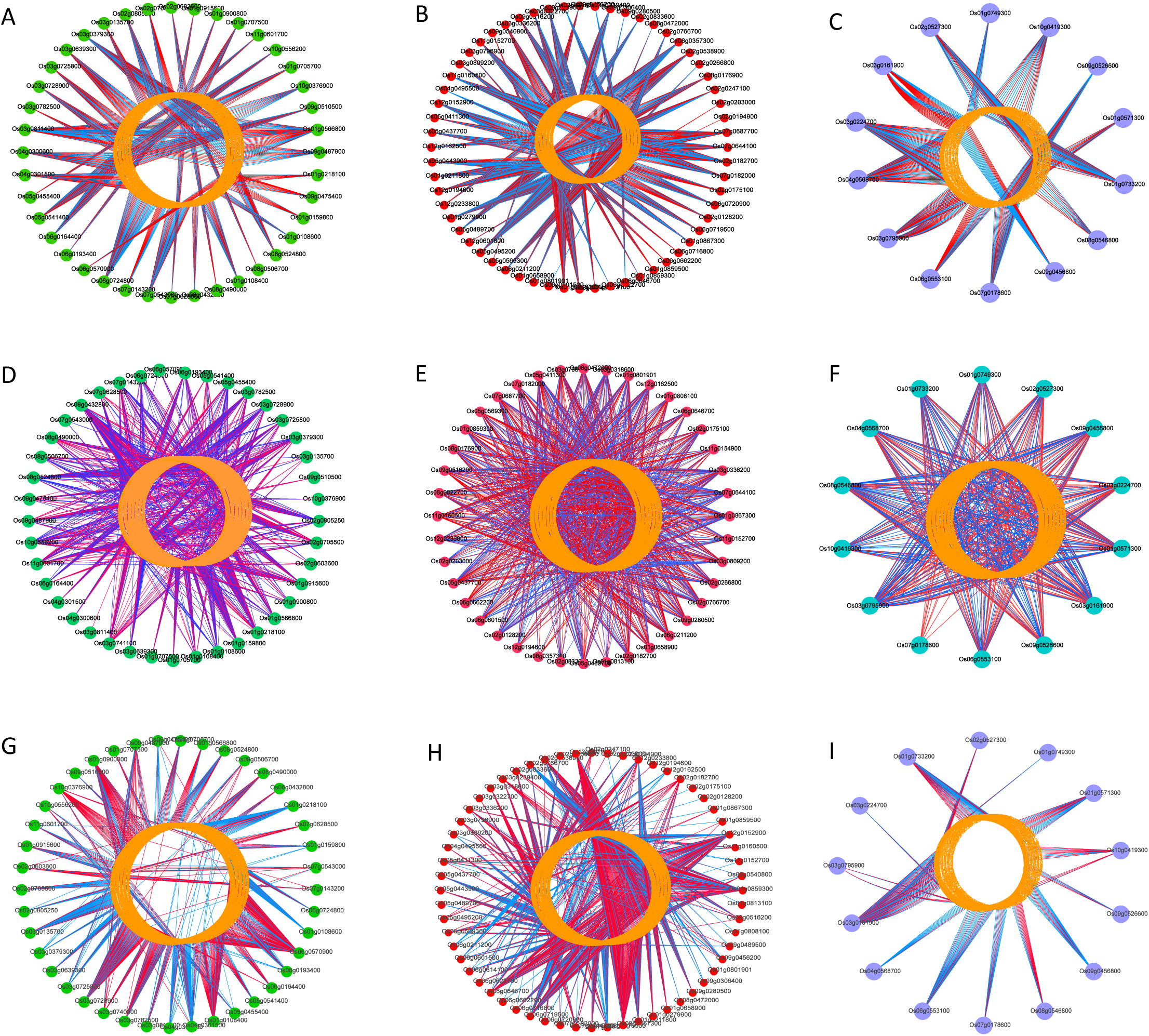
The correlation of TFs transcription level with the AS events PSI of downstream genes. The networks show the coefficient between the transcription levels of bHLH (left), bzip (middle) and hsfa (right) TF families and the AS events PSI of downstream genes under cold stress (A-C), drought stress (D-F) and flood stress (G-I).

### The effect of TF mutants on AS profiles of downstream genes

To further explore the effect of TFs on alternative splicing, we attempt to detect whether TF mutations can cause changes in splicing patterns of downstream genes, especially under environmental stresses. Taking into account of the availability of RNA-seq data for TF mutants under normal and stress conditions, we selected 3 TFs to compared download gene AS patterns between the mutants and wild lines. The results show extensive AS differences between mutants and wild lines. There are totally 535 and 719 DAS events between *bHLH148* mutant and wild lines under normal and drought condition, respectively (Figure 11A). 1272 and 1284 DAS events were detected between *bzip62* mutant and over-expressed lines with wild lines under drought stress (Figure 11B). 659 and 763 DAS events were found between *Oshsfa2e* mutant and wild lines under normal and drought condition, respectively (Figure 11C). DAS events between WT and mutants are significantly higher under stresses than normal conditions and the DAS events are mainly illustrated by DRI (Figure 11A) or DSE (Figure 11C), followed by DA3SS and DA5SS (Figure 11AC). We furether checked if the DASGs were overlapped with DEGs in TF mutants and the results show a small faction of genes were both DEGs and DASGs (Figure 11 D-F). Two representative examples were illustrated for a DA5SS event of gene *OsCPK4* between bzip62 over-expression lines and wild type (Figure 11G) and a DRI event of gene *OsDHSRP1* between bhlh148 mutant and wild type (Figure 11H). These results indicated that TFs can regulate different sets of downstream genes at AS or expression level, respectively and shed light on the new function of TFs on the regulation of AS.

**Figure 11.**
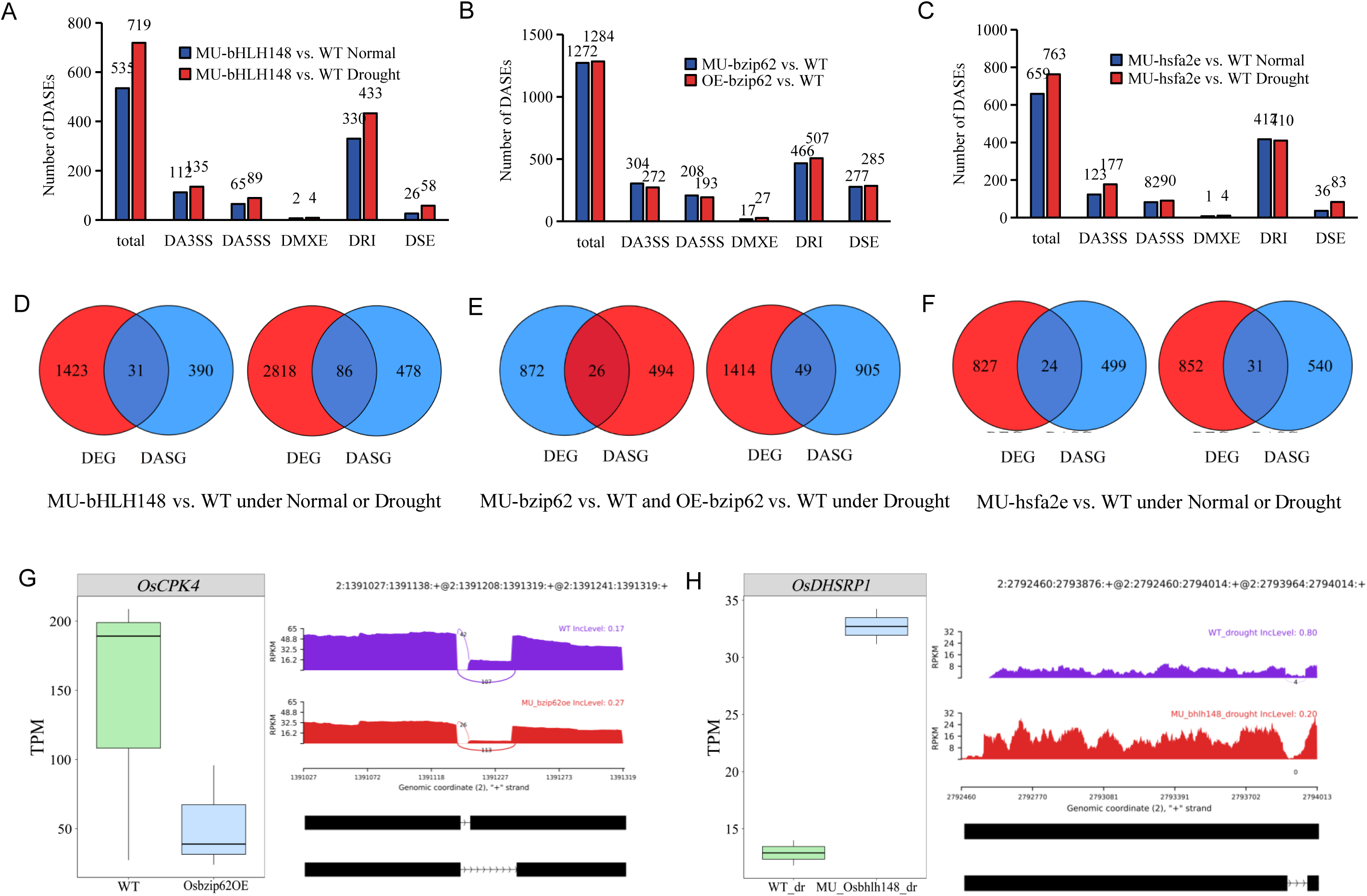
The effects of TFs on AS of downstream genes. (A-C) The number of total DAS and different types of AS events were shown for bHLH148 (A), bzip62 (B) and hsfa2e (C) TF mutants compared with wild-type lines. For A and C, normal and drought conditions were shown as blue and red, respectively. For B, the red and blue represent the comparison of mutant or overexpression lines with wild-type lines.

### Correlation analysis of transcription levels of TFs and SFs under abiotic stresses

In order to investigate the underlying mechanism underlying the effects of TFs on AS, we speculate that TF may have a regulatory effect on the transcription of SF factors, thus indirectly affecting AS of downstream genes. To validate this hypothesis, we analyzed the transcription levels correlation between TF and SF genes under stresses, and found the transcription levels of multiple TFs were significantly correlated with that of SFs (Figure 12). Among them, 27 bHLH family TFs were significantly correlated with the transcription levels of 14 SFs under drought treatments (Figure 12D). For example, *OsbHLH148* (Os03g0741100) significantly correlated with the transcriptional changes of SFs RS2Z38 and RS33. There are 40 BZIP TFs were significantly correlated with the expression changes of 15 SFs (Figure 12E) and 12 HSF transcription factors significantly correlated with the that of 9 SFs (Figure 12F). For example, *Oshsfa2e* (*Os03g0795900*) is significantly correlated with the expression changes of SR protein genes *RS2Z38*, *SR33a*, *RS2Z37*, *SCL26*, *SCL25*, *RSZ21a*, and *SCL30*. Similar results were shown for the transcription correlation analysis between TFs and SFs under other environmental conditions. Significantly, the closely correlation between expression changes of bHLH, bZIP and HSF TF families and SFs were demonstrated under cold stress (Figure 12 A-C) and water flooding conditions (Figure 12 G-I). These results indicate a correlation between the transcriptional changes of TFs and SFs under drought stress. We speculated that it is a widespread phenomena of TFs affecting AS by regulating the transcription of SFs under abiotic stresses. It is an additional regulation layer that TFs participating in the response to abiotic stresses by influencing SFs and then indirectly regulating alternative splicing.

**Figure 12.**
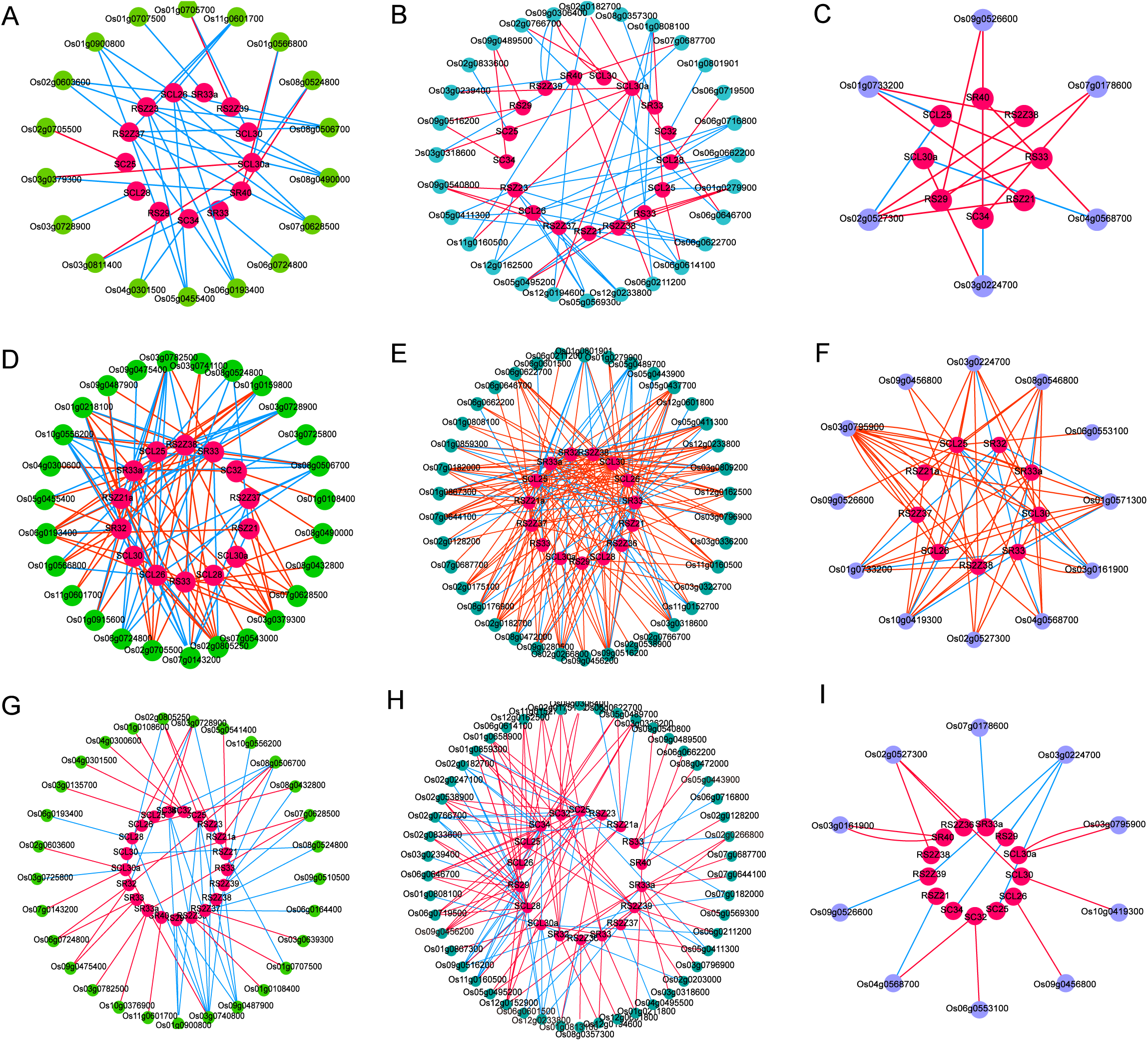
The correlation of TFs with SFs transcriptional level. The networks show the coefficient of transcription levels between bHLH (left), bzip (middle) and hsfa (right) TF families SFs under cold stress (A-C), drought stress (D-F) and flood stress (G-I).

**Figure 13.**
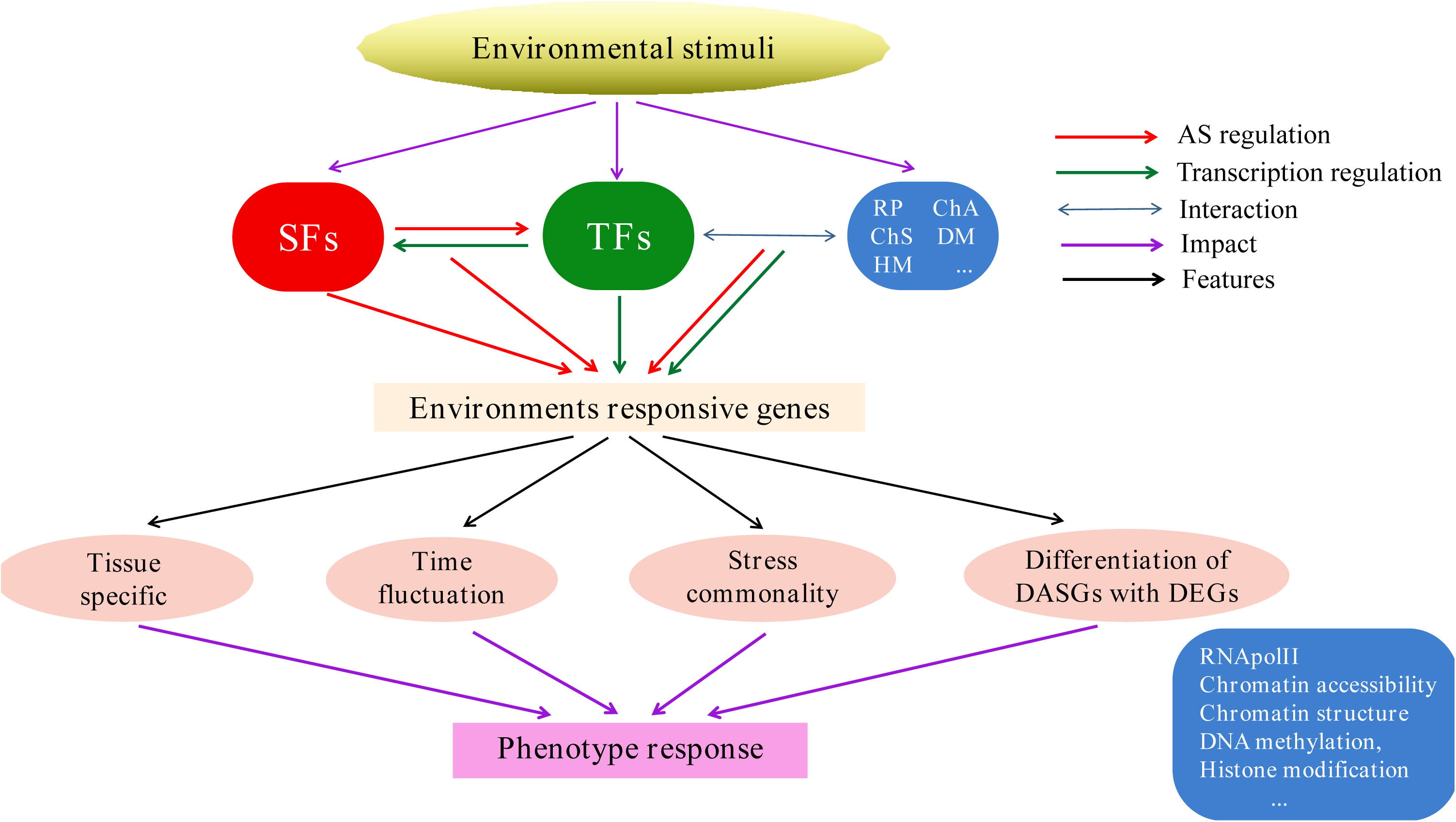
Model of universal AS features and AS regulation under diverse environmental stimuli. The regulation of the epigenetic elements and their interactions with TFs on AS were summarized from previous research or suggested for future research. Other features or regulation in the model were concluded from this study.

## Discussion

### AS Is widespread in rice under diverse environmental stimuli and shows universal characteristics

With global climate and environmental changes, plants are facing various environmental stresses such as drought, high salinity, extreme temperatures, heavy genus pollution and so on. AS is known to play essential roles in plant response to environmental signals. Traditionally, most of researches focus on changes in gene expression levels and ignore AS regulation. In this study, we found the important features of AS in response to environment stimuli, including AS tissue specificity, dynamic along treatment duration, commonality and differentiation among various environments, the less overlap between the response of AS and transcriptional level.

In detail, DASGs in response to environmental stresses show highly tissue specificity. The proportion of DASGs specific in roots or shoots is as high as 71-88% and their function enrichment are significantly different, except the common RNA binding or splicing related processes. For example, under low temperature stress, DASGs in shootss are mainly enriched RNA binding, starch biosynthesis processes, methyltransferase activity, helicase activity, MAPK cascade activation, mitochondrial transport, and RNA cleavage. Whereas in roots upon cold treatment, DASGs are mainly enriched in dephosphorylation, intracellular signal transduction, and abscisic acid activation signaling pathways, response damage, low temperature response and so on. Similar results were reported that splicing factors can be grouped as “basal” and “regulatory”, which are required to catalyze the splicing process itself and promote or repress splicing, respectively (Dvinge *et al*. 2019). The developmental stage-specific or tissue-specific or cell-type-specific variation in the expression level of regulatory splicing factors can confer a distinct splicing programs. Which indicates the tissue specific splicing program is a general phenomenon in both animal and plants. Therefore, in future research on AS responding to environmental signals, it is necessary to study more different tissues even cell types to obtain more rigorous rhythm.

We also found that the AS patterns of plants in response to environmental stress undergo dynamic changes over time, with a small fraction of genes continuously undergoing AS changes for a long period of time after stress. Whereas most genes only undergo AS changes in the early stages of stress occurrence, and subsequently exhibit AS pattern consistent with the control. This phenomena increases the complexity of plant response to stress. Further research is necessary to focus on the temporal dynamics of AS in response to environmental stimuli in order to accurately select candidate genes.

It is noteworthy that AS exhibits commonalities in response to different environmental conditions, with RNA splicing pathway related genes as prevalence. This is in line with earlier research. SR proteins were demonstrated to play critical roles in multiple mineral response, including Zn, Mn and P (Dong *et al*. 2018). Our results further highlight that AS of specific genes could not only be linked to single stress, but also to multiple various environmental stresses. In the future, it is necessary to conduct in-depth research on the functions of these common genes in regulating combined stresses in plants to explore the mechanisms of their pleiotropy and utilize them to improve plant adaptation to multi stresses.

Previous studies on plant responding to environmental stress have identified abundant DEGs, which play an important role in the regulation of plant response to environmental stress. In fact, pre-mRNA AS is also important regulation layers for plants in response to environmental changes. In this study we found only a small faction of overlap between DASGs and DEGs under multiple different external stimuli, and the functional categories enriched in DASGs and DEGs show few intersections. DASGs but not DEGs are mainly enriched in mRNA binding, splicing and so on, while DEGs but not DASGs are mainly relevant to ion transport, stress response, defense response regulation and signal transduction. Similar results were reported that AS is an additional mechanism beyond differential expression in rice under mineral nutrient treatments (Dong *et al*. 2018) and arabidopsis under termperature changes(James *et al*. 2012; Calixto *et al*. 2018). These results indicate that stress induced DEGs and DASGs affect different biological processes. It is necessary to conduct in-depth exploration of AS to decipher the comprehensive mechanisms by which plants respond to environmental stresses.

### TFs affect AS patterns under environmental stimuli through regulating the transcription of SFs

It is generally known that TFs play central roles in environmental response through regulating gene transcription level. In this study, we found that TFs functions not only by changing their own transcription level but also pre-mRNA splicing patterns. Briefly, some TFs themselves only show DE without DAS; some TFs show DAS but not DE; some TFs respond to environmental stimuli through both DE and DAS. The essential roles of AS changing in TF mRNAs might change the protein domains essential for DNA binding, dimerization, and transcriptional regulation (Seo *et al*. 2013). For example, AS of HsfA2 generating a small truncated HsfA2 isoform with an extra leucine-rich motif (Liu *et al*. 2013) and a maize NAC-domain retained splice variants interfering with the full-length NAC counterparts to confer Cd-tolerance function (Zhang *et al*. 2019). In this study, we found the generality that TFs can respond to diverse external stimuli by changing its transcription level as well as AS patterns.

Cummulating evidences indicated the regulation roles of TFs on AS (Malla *et al*. 2022; Ullah *et al*. 2023). We found the significant correlation between the transcription levels of some TFs and the PSI of stress response gene AS events, expecially the bHLH, bzip and hsfa TF families. We further provide direct evidence that the mutation or over-expression of three TFs, including *Osbhlh148,* Osbzip62 and *Oshsfa2e* can cause AS changes of download genes under stress. The similar research was reported in human that TFs are associated with AS events in the K562 cell line and the knockdown experiments of some TFs suggest their function as splicing enhancers (Ullah *et al*. 2023). In recent years, some TFs were found occurring at higher ration in intragenic regions than that in intergenic regions in both human genes and plants, suggesting a regulatory role of TFs beyond the regulation of gene expression (Han et al. 2017; Burgess et al. 2019).

Recently, emerging evidences revealed that TF can regulate the co-transcription process of AS. An important evidence in this field from human was strong over-representation of motifs relevant to some zinc finger TFs in IR events rather than non-retained introns (Ullah *et al*. 2023). Another demonstration about TFs involvment in co-transcriptional AS showed that TFs can regulate cell AS programs through controlling the expression of specific splicing regulators and regulate target AS programs via binding to RNA in human (Han et al. 2017). In plants, it was propose that hnRNP-like protein RZ-1C can promote efficient CTS through cooperative interaction with exons, introns and SFs (Ullah *et al*. 2023). However, the direct involvement of TFs in the regulation of AS in plants was rarely reported, especially comprehensive research in plants under diverse environmental stresses. Since SFs are well known regulatory factor for AS, in order to further explore the possible mechanism of TF regulating AS, we investigated the correlation between TFs and SFs. The results showed that multiple TFs co-expressed with SFs under diverse environmental stimuli. Therefore, we provide the direct evidence that TF was involved in the AS regulation and speculate that TFs might affect the splicing mode of stress responsive genes partly via regulating the transcription of SR gene in response to environmental stimuli, though the direct molecular mechanisms of TFs binding to mRNAs or physically regulating on the process of splicing were not demonstrated.

Notably, the epigenetic regulations on the co-transcriptional AS process has been emerging. Among of them, chromatin state play vital roles on exon skipping The regulatory functions of (Mercer et al. 2013) and intron retention (Ullah et al. 2018; Petrova et al. 2022). It is suggested that CHD1 can function as a chromatin remodeler indirectly affecting RNA splicing (Lee *et al*. 2017). The mechanisms conveying DNA methylation information into the regulation of alternative splicing was also illustrated (Lev Maor *et al*. 2015). In plants, a chromatin remodeler ZmCHB101 was demonstrated to impact AS contexts in response to osmotic stress (Petrova *et al*. 2022). With deep learning models, regions of open chromatin associated with IR from other intronic regions of open chromatin was distinguished (Ullah *et al*. 2018). However, the roles of transcription regulators such as TFs, chromatin remodeler on the co-transcriptional AS is still in the preliminary research stage, especially in plants responding to environmental stimuli. Further research needs to focus on the layers including the binding of TF to RNA, the relationship between TF and RNApolII extension, the synergistic regulation of AS by TF and chromatin accessibility, chromatin structure, DNA methylation, histone modification, and other functional mechanisms as well as exploring the splicing QTL to find important AS regulatory sites.

## Conclusion

Collectively, our work has revealed important general features of AS in response to environmental stimuli, the regulation roles of TFs on AS and the strong correlation of the transcription level of multiple TFs with SFs under various environmental conditions. We speculated that TFs might modulate the AS of download genes through regulating the transcription of SFs and discussed the recent advances and future research on co-transcriptional regulation of AS.

## Experimental procedures

### RNA-seq data

We selected datasets from TENOR (Transcriptome ENcyclopedia Of Rice, http://tenor.dna.affrc.go.jp) (Kawahara *et al*. 2016) and downloaded from the NCBI Sequence Read Archive (SRA; https://www.ncbi.nlm.nih.gov/sra; see Supplemental Table 1). A large scale of 238 mRNA-Seq libraries were provided from rice (Oryza. Sativa L. spp. japonica, var Nipponbare) under a wide variety of conditions, including abiotic stress conditions (high, low and extremely low cadmium; drought; osmotic; cold; and flood) and two plant hormone treatment conditions (ABA and jasmonic acid). The detailed samples preparation and growth conditions were described in previous study (Kawahara *et al*. 2016). Briefly, rice seedlings were treated with various Abiotic stress and Plant hormones at 10 days after germination (Table S1): (1) Cd stress: the seedlings were transferred to 1 µ M CdSO4, 0.2 µ M CdSO4 and 50 µ M CdSO4 for low, very low and high cadmium concentrations. Shoot and root tissues were taken from 0 h, 1, 4 and 10 days after low and extremely low cadmium treatment and 0 h, 12 h, and 5 days under high cadmium treatment; (2) Low temperature stress: shoots and roots were taken from 0 h, 1 h, 3 h, 6 h, 12 h, and 1 day after 4℃ cold treatment; (3) Flooding stress: completely immersed the seedlings in the culture medium and take shoots and roots from 0 h, 1 h, 3 h, 6 h, 12 h, 1 and 3 days after flooding treatment; (4) Osmotic stress: young shoots and roots were taken at 0 h, 1 h, 3 h, 6 h, 12 h after 0.6 M Mannitol treatment; (5) Drought stress: Take the seedlings from the culture medium and air dry them and take shoots and roots at 0 h, 1 h, 3 h, 6 h, 12 h, and 1 d after stress treatment; (6) Plant hormone treatment: transfer seedlings to 100 µ M JA or ABA medium and take young shoot and root tissue at 0 h, 1 h, 3 h, 6 h, 12 h and 1 day after treatment; (7) Normal growth and development: rice seedlings are not subject to any stress or hormone treatment. Young shoot and root tissues were taken after 0 h, 1 h, 3 h, 6 h, 12 h, 1, 3, 4, 5 and 10 days growth (Kawahara et al. 2016).

The RNA-seq dataset regarding to transcription factors mutants, over-expression plants and wild lines under various environmental conditions were also downloaded from the NCBI Sequence Read Archive (SRA; https://www.ncbi.nlm.nih.gov/sra; see Supplemental Table 1), including PRJNA272732 for *Oshsfa2e*, PRJNA272733 for *OsbHLH148*, PRJNA810084 for *HTG3* and PRJNA506820 for *OsbZIP62. OsHSFA2e*, *OsbHLH148* mutants and wild type rice plants (Oryza sativa ssp. japonica cv. Nipponbare) were exposed to controlled drought stress and well-watered conditions at the vegetative stage. Drought stress was applied on 45 day old plants and the soil water content was brought down to 40% field capacity over a period of 3-4 days and plants were maintained at that level for 10 days by weighing the pots daily at a fixed time of the day and replenishing the water lost through evapotranspiration. Another set of plants were maintained at 100% field capacity as well-watered (WW) condition (Gupta et al 2021). *HTG3* mutants and wild type plants grow until the four leaf stage and then were subjected to heat treatment at 42℃ for 0.5 hours or normal growth conditions for control (Wu *et al*. 2022). *OsbZIP62* mutants, over-expression and wild rice plants were grown normal conditions until four leaf stage (Yang *et al*. 2019).

### RNA-seq data pre-processing

An overview of the bioinformatic workflow in the present study is shown in Figure 1. In brief, FastQC was used to evaluate the sequencing quality and connectors. Trimmatic was applied to filter low quality and connectors to obtain high-quality Clean Data (Bolger *et al*. 2014). The genome sequence and annotation version Oryza_sativa. IRGSP-1.0 were downloaded from Ensembl Plants (http://plants.ensembl.org/). Hisat2 was applied to align Clean Reads to rice reference genome with default parameters (Kim et al 2015) and StringTie (Pertea et al 2015) was used to assemble each sample transcripts individually. TACO (Niknafs et al 2017) was used to compare and merge the individual GTF files with rice reference genome annotation files, resulting in a set of non redundant transcripts annotation. Afterwards, we use gffcompare (Pertea and Pertea 2020) to compare the transcript annotation file obtained in the previous step with the reference transcripts to classify, annotate, and evaluate the accuracy of the assembled transcript. Subsequently, a self-designed Python script was used to screen the results of gffcompare comparison (Dong et al. (2018)). The steps were as follows: (1) Extract transcripts with transcript classification codes of ‘j’, ‘o’, ‘u’, and ‘=’, which were used as candidate cleavage isomers detected in the dataset. ‘j’ represents transcripts that share at least one cleavage site with the reference transcript, ‘o’ represents the transcript with Exon overlapping with the reference transcript in the coding interval, ‘u’ represents the unknown intergenic transcript, which may be a new isomer from a known or new gene site, and ‘=’ represents the transcript that perfectly matches the reference transcript; (2) Retain transcripts with more than one Exon. Finally, a high-quality non redundant transcript annotated gtf file was obtained for subsequent analysis.

### Differential splicing and differential expression analysis

rMATS was used to identify alternative splicing events in each sample and differential AS events between treatments and normal conditions (Shen et al 2014). The conditions for screening differential AS events are: 1) The sum of all biological repeat IJC (Inclusion junction count) in each sample is greater than or equal to 1, and the sum of SJC (Skipping junction count) is greater than or equal to 1. Take the exon skipping events as an example, IJC represents the number of reads covered by the splicing site at the junction of both sides of the Exon and the middle Exon, and SJC represents the number of reads covered by the splicing site at the junction of both sides of the Exon; 2) IJC+SJC ≥ 5x, FDR ≤ 0.05, and ΔPSI ≥ 0.1, where x is the sum of biological replicates in the control and treatment groups, and PSI (Percentage spliced in) is the percentage of cleavage, ΔPSI is the difference in PSI between the treatment and the control, calculated as PSI=(IJC/IncFormLen)/(IJC/IncFormLen)+(SJC/SkipFormLen), ΔPSI=|PSI(control)-PSI(treatment) |.

FeatureCounts was used to calculate the reads covered on genes and DESeq2 was used to analyze the difference between the treatment and control samples (Liao et al 2014) (Love et al 2014). The value of gene expression was represented as TPM (Transcripts Per Million). The screening conditions of differentially expressed genes were |log2FoldChange| > 1 and Pvalue < 0.01.

### Annotation and functional enrichment analysis

GO and KEGG enrichment analysis were conducted in Plant Regulomics (http://bioinfo.sibs.ac.cn/plant-regulomics/) (Ran et al 2020). Oryza sativa (RAP-DB) was selected for the species option and a list of differentially expression or AS genes were submitted for GO and KEGG enrichment analysis, with the significant enrichment threshold P<0.05. Subsequently, R package ComplexHetmap (Gu et al 2016) was used to visualize the annotation results.

### Differentially expression and differentially AS analysis of SFs and TFs during various environmental stimuli

22 splicing related proteins in six SR protein subfamilies were identified in rice (Barta et al. (2010)). We selected these 22 SR proteins as the representatives of splicing factors to evaluate their regulation function on alternative splicing and subsequent association analysis with transcription factors. We extracted the gene expression and splicing patterns of these 22 SRs at each time point under various conditions, to comprehensively analyze the gene expression and splicing mode of SR factors under abiotic stress and plant hormone treatments. Annotated Information of TFs was downloaded from PlantTFDB (http://planttfdb.gao-lab.org/). Subsequently, we screened out the transcription factors that are simultaneously differentially expressed and differentially alternative spliced under various abiotic stresses or hormone treatments. HeatMap function in TBtools (Chen et al 2020) was utilized for visualization.

### Correlation analysis of TFs with splicing ration (PSI) and network construction

To investigate the possible regulation relationship between TFs and splicing events, we calculated the spearman correlation coefficient of expression level of TFs and PSI value of splicing events under each time point under various treatments with cor.test function in R. Cytoscape was used to visualize the relationship between TFs and splicing events (Shannon et al. 2003).

### Gene expression and Alternative splicing analysis in SFs and TFs Mutants

Gene expression and Alternative splicing analysis in SFs and TFs Mutants was conducted with the same method as that use in multi time durations under different abiotic stresses. The DEGs and DASGs analysis were compared between mutants and wild type under normal and stress conditions. Especially, the DE or DAS SR coding genes upon TFs mutation were analyzed under normal and stress treatments. Pheatmap in R was used to visualize the expression and splicing patterns. To investigate the possible regulation roles of TFs on SRs, we further analyzed the relationship of TFs and SFs, calculated as the spearman correlation coefficient of expression level of SRs and TFs at different durations of drought stress (r>0.95, p<0.05). The significant relationship was visualized with Cytoscape 3.7.1.

## Legend

Figure S1 The distribution of intron number (A) and intron length (B).

Figure S2 Overview of alternative splicing identified in rice shoot and root under various growth conditions.

Figure S3 Functional GO analysis of DASGs in rice shoots and roots. Graphs show functional categorization (GO) of DASGs in shoots (blue) and roots (red) under cadmium (A), ABA (B) or JA (C) growth conditions. The log10(1/P) values represents the enrichment ratio of each GO terms.

Figure S4 The temporal fluctuation of stress-induced DASGs. The time specific and shared DASGs are shown for shoots (A) and roots (B) under flood stress or for shoots (C) and roots (D) under osmotic stress.

Figure S5 The temporal fluctuation of stress-induced DASGs. The DASGs were dynamically changing with the treatment duration under environmental stimuli. The time specific and shared DASGs are shown for shoots (A) and roots (D) under very low Cd stress or for shoots (B) and roots (E) under low Cd stress or for shoots (C) and roots (F) unde high Cd stress.

Figure S6 The temporal fluctuation of stress-induced DASGs. The DASGs were dynamically changing with the treatment duration under environmental stimuli. The time specific and shared DASGs are shown for shoots (A) and roots (B) under JA treatment or for shoots (C) and roots (D) under ABA treatment.

Figure S7 The enrichment GO terms for shared DASGs under different concentration of Cd stress were shown for shoots (A) and roots (B) and under ABA and JA treatments were shown for shoots (C) and roots (D).

Figure S8 Comparison of DEGs and DASGs under flood or osmotic stress. (A-D) The line charts and venn diagrams represent the number and overlap of DEGs, ASGs and DASGs in shoots and roots under different flood and osmotic stress.

Figure S9 Comparison of DEGs and DASGs under Cd stress. (A-F) The line charts and venn diagrams represent the number and overlap of DEGs, ASGs and DASGs in shoots and roots under very low, low or high concentration of Cd stress.

Figure S10 Comparison of DEGs and DASGs under hormone treatments. (A-D) The line charts and venn diagrams represent the number and overlap of DEGs, ASGs and DASGs in shoots and roots under different JA and ABA treatment durations.

Figure S11 Comparison of GO annotation of DEGs and DASGs under Cd stress. (A-F) The enrichment GO terms for DASGs (red) and DEGs (blue) in shoots and roots under very low, low or high concentration of Cd stress.

Figure S12 Comparison of GO annotation of DEGs and DASGs under hormone treatments. (A-D) The enrichment GO terms for DASGs (red) and DEGs (blue) in shoots and roots under ABA or JA treatment.

Table S1 RNA-seq data used in this study

Table S2-15 DAS events information in each tissue under every condition

Table S16 The transcription factors showing DEG and DASG simultaneously in each condition

## Acknowledgments

This work was supported by the self-determined research fund of Central China Normal University (CCNU18QN027), the National Special Key Project of China on Transgenic Research (2016ZX 08001-003), National Natural Science Foundation of China (31371550) and National Key Research & Development Program (2017YFD0300106).

## Author contributions

Z.Z. and B.X. designed the research; B.X, S.Y., C.W., F.Z., Y.L. Z.X. G.X. and Z.Z. analyzed data; and B.X. and Z.Z. wrote the manuscript. X.G. and Z.Z. revised the manuscript. Z.Z. finalized the manuscript. All authors agreed the final version of the manuscript.

## Reference

A A., H B., Id O., GK K., I B. & MM M. (2023) - SCR106 splicing factor modulates abiotic stress responses by maintaining the RNA splicing in rice. LID - erad433 [pii] LID - 10.1093/jxb/erad433 [doi]. J Exp Bot 31.

Bolger A., Lohse M. & Usadel B. (2014) Trimmomatic: a flexible trimmer for Illumina sequence data. Bioinformatics 30, 2114–20.

Calixto C., Guo W., James A., Tzioutziou N., Entizne J., Panter P., Knight H., Nimmo H., Zhang R. & Brown J. (2018) Rapid and Dynamic Alternative Splicing Impacts the Arabidopsis Cold Response Transcriptome. Plant Cell 30, 1424–44.

Chen Y., Yang W., Zou Y., Wu Y., Mao W., Zhang J., Zia-ur-Rehman M., Wang B. & Wu P. (2024) Quantification of the effect of biochar application on heavy metals in paddy systems: Impact, mechanisms and future prospects. Science of The Total Environment 912, 168874.

Dong C., He F., Berkowitz O., Liu J., Cao P., Tang M., Shi H., Wang W., Li Q., Shen Z., Whelan J. & Zheng L. (2018) Alternative Splicing Plays a Critical Role in Maintaining Mineral Nutrient Homeostasis in Rice (Oryza sativa). Plant Cell 30, 2267–85.

Dvinge H., Guenthoer J., Porter P. & Bradley R. (2019) - RNA components of the spliceosome regulate tissue- and cancer-specific alternative splicing. Genome Res 29, 1591–604.

Eckardt N.A., Ainsworth E.A., Bahuguna R.N., Broadley M.R., Busch W., Carpita N.C., Castrillo G., Chory J., DeHaan L.R., Duarte C.M., Henry A., Jagadish S.V.K., Langdale J.A., Leakey A.D.B., Liao J.C., Lu K.J., McCann M.C., McKay J.K., Odeny D.A., Jorge de Oliveira E., Platten J.D., Rabbi I., Rim E.Y., Ronald P.C., Salt D.E., Shigenaga A.M., Wang E., Wolfe M. & Zhang X. (2023) Climate change challenges, plant science solutions. Plant Cell 35, 24–66.

Han H., Braunschweig U., Gonatopoulos-Pournatzis T., Weatheritt R., Hirsch C., Ha K., Radovani E., Nabeel-Shah S., Sterne-Weiler T., Wang J., O’ Hanlon D., Pan Q., Ray D., Zheng H., Vizeacoumar F., Datti A., Magomedova L., Cummins C., Hughes T., Greenblatt J., Wrana J., Moffat J. & Blencowe B. (2017) - Multilayered Control of Alternative Splicing Regulatory Networks by Transcription Factors. Mol Cell 65, 539–53.

Jabre I., Reddy A.S.N., Kalyna M., Chaudhary S., Khokhar W., Byrne L.J., Wilson C.M. & Syed N.H. (2019) Does co-transcriptional regulation of alternative splicing mediate plant stress responses? Nucleic Acids Res 47(6), 11.

James A.B., Syed N.H., Bordage S., Marshall J., Nimmo G.A., Jenkins G.I., Herzyk P., Brown J.W. & Nimmo H.G. (2012) Alternative splicing mediates responses of the Arabidopsis circadian clock to temperature changes. Plant Cell 24, 961–81.

Kawahara Y., Oono Y., Wakimoto H., Ogata J., Kanamori H., Sasaki H., Mori S., Matsumoto T. & Itoh T. (2016) TENOR: Database for Comprehensive mRNA-Seq Experiments in Rice. Plant Cell Physiol 57, e7.

Lee Y., Park D. & Iyer V.R. (2017) The ATP-dependent chromatin remodeler Chd1 is recruited by transcription elongation factors and maintains H3K4me3/H3K36me3 domains at actively transcribed and spliced genes. Nucleic Acids Res 45, 7180–90.

Lev Maor G., Yearim A. & Ast G. (2015) The alternative role of DNA methylation in splicing regulation. Trends Genet 31, 274–80.

Li J., Zhang Z., Chong K. & Xu Y. (2022) Chilling tolerance in rice: Past and present. Journal of Plant Physiology 268, 153576.

Li X.L., Meng, D., Li, M. J., Zhou, J., Yang, Y. Z., Zhou, B. B., Wei, Q. P., & Zhang, J. K. (2023) Transcription factors MhDREB2A/MhZAT10 play a role in drought and cold stress response crosstalk in apple.

Ling Y., Mahfouz M.M. & Zhou S. (2021) Pre-mRNA alternative splicing as a modulator for heat stress response in plants. Trends in Plant Science 26, 1153–70.

Liu J., Sun N., Liu M., Du B., Wang X. & Qi X. (2013) An autoregulatory loop controlling Arabidopsis HsfA2 expression: role of heat shock-induced alternative splicing. Plant Physiol 162, 512–21.

Malla S., Prasad Bhattarai D., Groza P., Melguizo-Sanchis D., Atanasoai I., Martinez-Gamero C., Roman A.C., Zhu D., Lee D.F., Kutter C. & Aguilo F. (2022) ZFP207 sustains pluripotency by coordinating OCT4 stability, alternative splicing and RNA export. EMBO Rep 23, e53191.

Mercer T.R., Edwards S.L., Clark M.B., Neph S.J., Wang H., Stergachis A.B., John S., Sandstrom R., Li G., Sandhu K.S., Ruan Y., Nielsen L.K., Mattick J.S. & Stamatoyannopoulos J.A. (2013) DNase I-hypersensitive exons colocalize with promoters and distal regulatory elements. Nat Genet 45, 852–9.

Mingde Deng Y.W., Monika Kuzma, Maryse Chalifoux, Linda Tremblay, Shujun Yang, Jifeng Ying, Angela Sample, Hung-Mei Wang, Rebecca Griffiths, Tina Uchacz, Xurong Tang, Gang Tian, Katelyn Joslin, David Dennis, Peter McCourt, Yafan Huang, Jiangxin Wan (2023) Transcription factors MhDREB2A/MhZAT10 play a role in drought and cold stress response crosstalk in apple. Plant Physiol 192, 2203–20.

Petrillo E. (2023) Do not panic: An intron-centric guide to alternative splicing. Plant Cell 35, 1752–61.

Petrova V., Song R., Nordstrom K.J.V., Walter J., Wong J.J.L., Armstrong N.J., Rasko J.E.J. & Schmitz U. (2022) Increased chromatin accessibility facilitates intron retention in specific cell differentiation states. Nucleic Acids Res 50, 11563–79.

Reddy A.S., Marquez Y Fau - Kalyna M., Kalyna M Fau - Barta A. & Barta A. Complexity of the alternative splicing landscape in plants.

Renard D. & Tilman D. (2019) National food production stabilized by crop diversity. Nature 571, 257–60.

Sarma B., Kashtoh, H., Lama Tamang, T., Bhattacharyya, P. N., Mohanta, Y. K., & Baek, K. H. (2023) Abiotic Stress in Rice: Visiting the Physiological Response and Its Tolerance Mechanisms. LID - 10.3390/plants12233948 [doi] LID - 3948. Plants 12.

Saud S., Wang D., Fahad S., Alharby H.F., Bamagoos A.A., Mjrashi A., Alabdallah N.M., AlZahrani S.S., AbdElgawad H., Adnan M., Sayyed R.Z., Ali S. & Hassan S. (2022) Comprehensive Impacts of Climate Change on Rice Production and Adaptive Strategies in China. Front Microbiol 13, 926059.

Seo P.J., Park M.J. & Park C.M. (2013) Alternative splicing of transcription factors in plant responses to low temperature stress: mechanisms and functions. Planta 237, 1415–24.

Shahzad A., Ullah S., Dar A.A., Sardar M.F., Mehmood T., Tufail M.A., Shakoor A. & Haris M. (2021) Nexus on climate change: agriculture and possible solution to cope future climate change stresses. Environ Sci Pollut Res Int 28, 14211–32.

Ullah F., Hamilton M., Reddy A.S.N. & Ben-Hur A. (2018) Exploring the relationship between intron retention and chromatin accessibility in plants. BMC Genomics 19, 21.

Ullah F., Jabeen S., Salton M., Reddy A.S.N. & Ben-Hur A. (2023) Evidence for the role of transcription factors in the co-transcriptional regulation of intron retention. Genome Biol 24, 53.

Wu N., Yao Y., Xiang D., Du H., Geng Z., Yang W., Li X., Xie T., Dong F. & Xiong L. (2022) - A MITE variation-associated heat-inducible isoform of a heat-shock factor confers heat tolerance through regulation of JASMONATE ZIM-DOMAIN genes in rice. New Phytol 234, 1315–31.

X Y., Z J., Q P., Y T., F Z., Id O. & Y L. (2022) - ABA Mediates Plant Development and Abiotic Stress via Alternative Splicing. LID - 10.3390/ijms23073796 [doi] LID - 3796. Int J Mol Sci 23.

Yang S., Xu K., Chen S., Li T., Xia H., Chen L., Liu H. & Luo L. (2019) A stress-responsive bZIP transcription factor OsbZIP62 improves drought and oxidative tolerance in rice. BMC Plant Biology 19, 260.

Yang X., Jia Z., Pu Q., Tian Y., Zhu F. & Liu Y. (2022) ABA Mediates Plant Development and Abiotic Stress via Alternative Splicing. Int J Mol Sci 23.

Ye L.X., Wu Y.M., Zhang J.X., Zhou H., Zeng R.F., Zheng W.X., Qiu M.Q., Zhou J.J., Xie Z.Z., Hu C.G. & Zhang J.Z. (2023) A bZIP transcription factor (CiFD) regulates drought- and low-temperature-induced flowering by alternative splicing in citrus. J Integr Plant Biol 65, 674–91.

Zhang J., Li L., Huang L., Zhang M., Chen Z., Zheng Q., Zhao H., Chen X., Jiang M. & Tan M. (2019) Maize NAC-domain retained splice variants act as dominant negatives to interfere with the full-length NAC counterparts. Plant Sci 289, 110256.

Zhang Z. & Xiao B. (2018) Comparative alternative splicing analysis of two contrasting rice cultivars under drought stress and association of differential splicing genes with drought response QTLs. Euphytica 214, 73.

